# Testosterone-mediated behavior shapes the emergent properties of social networks

**DOI:** 10.1101/737650

**Authors:** Roslyn Dakin, Ignacio T. Moore, Brent M. Horton, Ben J. Vernasco, T. Brandt Ryder

**Affiliations:** Migratory Bird Center, Smithsonian Conservation Biology Institute, National Zoological Park, Washington, DC 20013, USA; Department of Biology, Carleton University, Ottawa, Ontario K1S 5B6, Canada; Department of Biological Sciences, Virginia Tech, Blacksburg, Virginia 24061, USA; Department of Biology, Millersville University, Millersville, Pennsylvania 17551, USA; Bird Conservancy of the Rockies, Fort Collins, Colorado, 80525, USA

**Keywords:** androgens, behavioral endocrinology, collective behavior, cooperation, dynamic networks, social networks, testosterone

## Abstract

1. Social networks can vary in their organization and dynamics, with implications for ecological and evolutionary processes. Understanding the mechanisms that drive social network dynamics requires integrating individual-level biology with comparisons across multiple social networks.
2. Testosterone is a key mediator of vertebrate social behavior and can influence how individuals interact with social partners. Although the effects of testosterone on individual behavior are well established, no study has examined whether hormone-mediated behavior can scale up to shape the emergent properties of social networks.
3. We investigated the relationship between testosterone and social network dynamics in the wire-tailed manakin, a lekking bird species in which male-male social interactions form complex social networks. We used an automated proximity system to longitudinally monitor several leks and we quantified the social network structure at each lek. Our analysis examines three emergent properties of the networks: social specialization (the extent to which a network is partitioned into exclusive partnerships), network stability (the overall persistence of partnerships through time), and behavioral assortment (the tendency for like to associate with like). All three properties are expected to promote the evolution of cooperation. As the predictor, we analyzed the collective testosterone of males within each social network.
4. Social networks that were composed of high-testosterone dominant males were less specialized, less stable, and had more negative behavioral assortment, after accounting for other factors. These results support our main hypothesis that individual-level hormone physiology can predict group-level network dynamics. We also observed that larger leks with more interacting individuals had more positive behavioral assortment, suggesting that small groups may constrain the processes of homophily and behavior-matching.
5. Overall, these results provide evidence that hormone-mediated behavior can shape the broader architecture of social groups. Groups with high average testosterone exhibit social network properties that are predicted to impede the evolution of cooperation.

## Introduction

Behavioral interactions are the foundation of social network structures that can vary through time, among populations, and across species. Network structures play an important role in many ecological and evolutionary processes, including the spread of diseases (Sah *et al*. 2018; Stroeymeyt *et al*. 2018), the transmission of information and resources (Aplin *et al*. 2012; Maldonado-Chaparro *et al*. 2018), and selection on individual behavior (Ohtsuki *et al*. 2006). A major challenge is understanding how individual-level factors, such as physiological and behavioral mechanisms, scale up to drive the emergent structural properties of social groups (Krause & Ruxton 2002). Linking these two levels of analysis is difficult because it requires integrating individual-level data with repeated measures of entire social groups (Sah *et al*. 2017).

Here, we use a comparison of multiple social networks through time to investigate how hormone-mediated behavior shapes the higher-order structure of social networks. Testosterone is a steroid hormone that is well known for its influence on social behavior and its sensitivity to changes in the social environment (Wingfield *et al*. 1990; Adkins-Regan 2005; Goymann 2009). Testosterone often promotes physical aggression and other behaviors associated with social dominance (Oyegbile & Marler 2005; Fuxjager *et al*. 2010). Testosterone can also promote status-seeking behaviors in a non-aggressive context, including cooperative and gregarious behavior (Eisenegger *et al*. 2011; Boksem *et al*. 2013; Ryder *et al*. 2020). Overall, these hormone-signaling pathways are essential for the development and modulation of complex behavioral phenotypes (Cohen *et al*. 2012).

To investigate how testosterone is associated with social network dynamics, we studied the wire-tailed manakin (*Pipra filicauda*), a bird species in which the males engage in coordinated displays with each other at sites known as leks. In *P. filicauda*, males exhibit two social status classes: the dominant males who hold territories on the leks, and subordinate, “floater” males who must acquire a territory on a lek before they can mate (Heindl 2002; Ryder *et al*. 2011a). Male social partnerships in wire-tailed manakins can be remarkably stable and typically occur between two unrelated males, most commonly, but not exclusively, between a territory-owner and a floater (Ryder *et al*. 2011a; Dakin & Ryder 2020). Two features of the manakin leks make them especially well-suited to studying the relationship between hormones and group-level social structure. First, testosterone is known to affect the social behavior of individual male wire-tailed manakins (Ryder *et al*. 2020). Second, these partnerships among males form the basis of complex and dynamic social networks that are replicated across leks, facilitating a comparative approach (Dakin & Ryder 2018, 2020). Given this background, we sought to test whether testosterone and its effects on male behavioral phenotype could drive the emergent properties of the social network.

The broader function of male-male social behavior and coordinated displays in the manakin family has been the subject of considerable study (e.g., Prum 1994; DuVal 2007; McDonald 2007; Ryder *et al*. 2008, 2009; Díaz-Muñoz *et al*. 2014). One possible function is that dyadic displays may be competitive and/or they may serve to maintain an individual’s position in a dominance hierarchy (Prum 1994; Heindl 2002). Social interactions may also represent cooperative coalitions that provide benefits to both parties (Ryder *et al*. 2008, 2009). The potential competitive and cooperative functions of male-male social behavior are not mutually exclusive, and function may be context- and/or status-specific (Ryder *et al*. 2008). Coordinated displays may also be a vestige of ancestral cooperative behavior (i.e., the behavior may have been directly beneficial to both parties in the past, and it persists today, even if it no longer has adaptive benefits; Prum 1994). Given this background and recent evidence that testosterone modulates male social behavior in wire-tailed manakins (Ryder *et al*. 2020), we focused this study on three emergent properties of the social networks formed by male-male interactions that can influence the evolution and maintenance of cooperation (Fig. 1).

**Figure 1.**
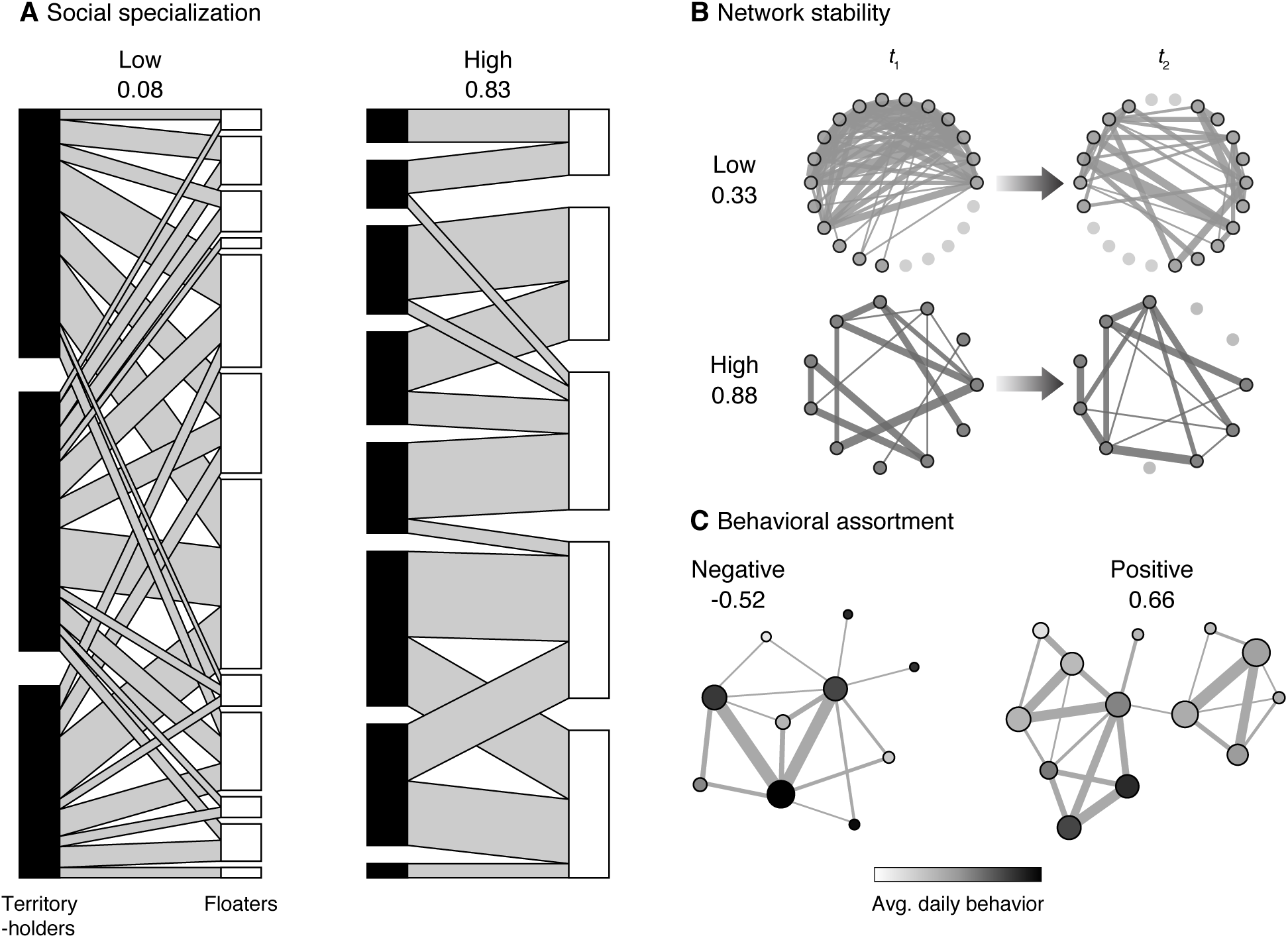
Example social networks illustrating social specialization, network stability, and behavioral assortment. (A) Social specialization is measured using the bipartite form of the social network. In the example on the left, the associations between floater males and territory-holders are poorly partitioned, creating a network with a relatively low specialization. On the right, there is greater partitioning, such that each floater male maintains a smaller number of associations with specific territory-holding males. (B) Network stability is measured by evaluating the persistence of partnerships from one recording session (t_1_) to the next (t_2_). The example at the top of (B) has a lower stability than the example on the bottom. (C) Behavioral assortment measures the tendency of like to associate with like. In (C), nodes are shaded to indicate a continuous gradient of daily behavior (in this case, each male’s average daily strength is shown). In the example network on the left, the males tend to associate with behaviorally dissimilar partners, yielding a negative behavioral assortment. In the example on the right, the males tend to associate with behaviorally similar partners, yielding a positive assortment. See Table 1 for additional data.

The first property we examined, social specialization, seeks to capture the exclusivity of the relationships between partners in a social network. In the context of manakin behavior, highly specialized networks are well-partitioned among specific territory-owner and floater relationships, as illustrated in Fig. 1A. In humans, social specialization has been found to maximize the ability of a team to successfully perform a challenging task (Jehn & Shah 1997). In manakins, we expect specialization to improve the familiarity of social partners and the behavioral coordination of their displays. Greater specialization is also expected to minimize conflict over mating and territorial ascension opportunities (Schjelderup-Ebbe 1922; McDonald 1993).

**Table 1.** Descriptive statistics for manakin social networks. Means and standard deviations (SD) are provided for each field season. The bottom row provides the sample sizes for the number of social networks analyzed (n_obs_) at each lek (N_lek_). Additional data is provided in the supplement in Fig. S1 and Fig. S4.

| Field season | Network size<br>(# birds) | Social<br>specialization | Network<br>stability | Behavioral<br>assortment | Collective T<br>(terr.) | Collective T<br>(floa.) |
| --- | --- | --- | --- | --- | --- | --- |
| 15-16 | 14.5 (8.9) | 0.50 (0.22) | 0.43 (0.25) | 0.01 (0.36) | 0.44 (0.42) | -0.31 (0.56) |
| 16-17 | 17.8 (10.5) | 0.30 (0.24) | 0.47 (0.20) | 0.06 (0.30) | 0.49 (0.24) | -0.22 (0.37) |
| 17-18 | 10.3 (4.5) | 0.42 (0.30) | 0.42 (0.27) | 0.02 (0.37) | 0.43 (0.40) | -0.70 (0.46) |
| Sample size | $n_{\text{obs}} = 80$ | $n_{\text{obs}} = 70$ | $n_{\text{obs}} = 54$ | $n_{\text{obs}} = 80$ | $n_{\text{obs}} = 80$ | $n_{\text{obs}} = 80$ |
| | $N_{\text{lek}} = 11$ | $N_{\text{lek}} = 11$ | $N_{\text{lek}} = 11$ | $N_{\text{lek}} = 11$ | $N_{\text{lek}} = 11$ | $N_{\text{lek}} = 11$ |

The second property, network stability (Fig. 1B), quantifies the average persistence of social partnerships through time (Poisot *et al*. 2012; Dakin & Ryder 2020). Coordinated displays in manakins require the synchronization of complex behaviors, and previous empirical work indicates that longer partnership tenure has a positive effect on display coordination (Trainer & McDonald 1995; Trainer *et al*. 2002). Greater temporal stability of partnerships also increases the opportunity for familiarity and reciprocity within a social network (Trivers 1971; Roberts & Sherratt 1998; Croft *et al*. 2006).

The third property, behavioral assortment (Fig. 1C), captures the extent to which males interact with partners who express similar behaviors (i.e., is like associated with like? Croft *et al*. 2006; Farine 2014). At the proximate level, positive assortment may represent the outcome of generalized reciprocity and/or partner choice (Fowler & Christakis 2010; Dakin & Ryder 2018). At the ultimate level, positive assortment has also been shown to promote the evolution of cooperation (Ohtsuki *et al*. 2006). To quantify the overall behavioral assortment of the manakin networks, we focused on two correlated metrics of social behavior within the network: “strength” (a male’s frequency of daily social interactions) and “degree” (his daily number of social partnerships; Dakin & Ryder 2018). We used a composite measure of assortment that averaged the assortativity indices of these two phenotypes.

As a potential predictor of specialization, stability, and assortment, we quantified the collective testosterone of the manakin leks (Fig. 2; Akinola *et al*. 2016). Because hormone-behavior relationships are status-specific in manakins and many other species (Eisenegger *et al*. 2011; Boksem *et al*. 2013; Ryder *et al*. 2020), we analyzed collective testosterone for each of the two status classes separately. High individual testosterone levels are associated with reduced sociality in dominant males, but increased sociality in subordinate males (Ryder *et al*. 2020). We therefore predicted that the average (or collective) testosterone of territorial males in a group would be negatively associated with its specialization, stability, and assortment. In contrast, we predicted that the collective testosterone of floater males in a group would be positively associated with these three emergent properties of the social network.

**Figure 2.**
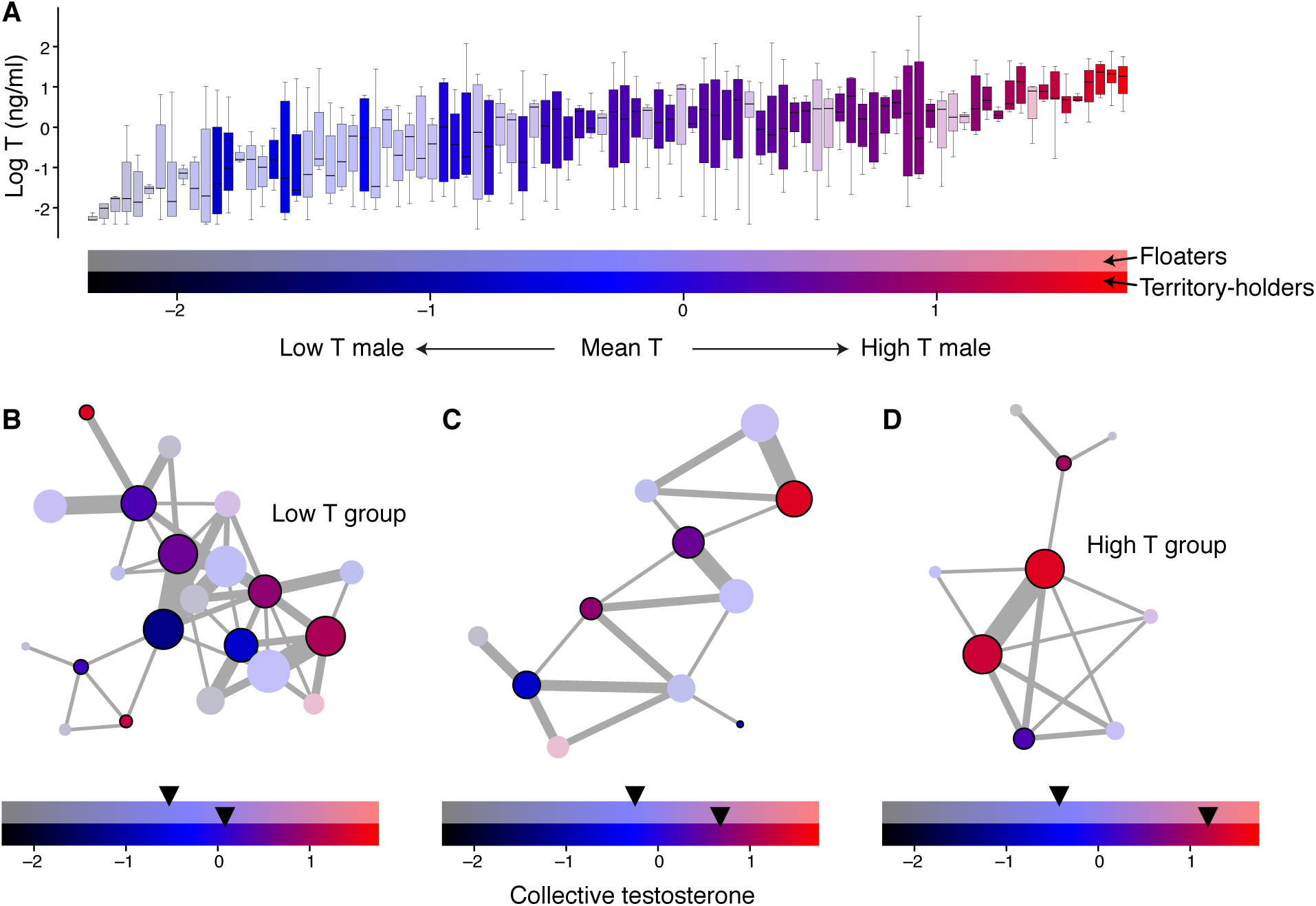
Collective testosterone of manakin social networks. (A) Circulating testosterone varies among individual males. This graph shows repeated measures from 210 individual male manakins, sorted along the x-axis by mean testosterone, a standardized measure of a male’s average circulating testosterone level (Ryder *et al*. 2020). Opacity is used to denote the two male status classes (with subordinate floaters colored semi-transparent, and dominant territory-holders colored opaque). Color ramping is used to denote each individual’s hormone phenotype. (B-D) These data were used to define collective testosterone in the present study. Node colors indicate an individual’s mean testosterone following the scale in (A). The collective testosterone of each social network is calculated as the average of the individual hormone phenotypes, weighted by strength, and is shown below each example. Collective testosterone was determined for each status class separately. See Table 1 for additional data.

## Materials and Methods

### Study Population

We studied wire-tailed manakins (*Pipra filicauda*) at the Tiputini Biodiversity Station in Orellana Province, Ecuador (0° 38’ S, 76° 08’ W). This population of *P. filicauda* has been studied and individuals color-banded annually since 2002 (e.g., Ryder *et al*. 2008, 2009). The present study was conducted on 11 leks during peak breeding activity (December-March) across three field seasons: 2015-16, 2016-17 and 2017-18. All research was approved by the Smithsonian ACUC (protocols #12-23, 14-25, and 17-11) and the Ecuadorean Ministry of the Environment (MAE-DNB-CM-2015-0008).

### Testosterone Assessment

Male manakins were captured using mist-nets on the leks as described in (Ryder *et al*. 2020). We deployed up to 16 mist-nets simultaneously at a given lek, with the intention of capturing every male on the lek. We rotated the nets between leks with the goal of capturing each male up to three times per field season. Each mist-net was checked on a 30-minute schedule with variation resulting from capture rate (i.e., multiple birds being caught on the same net run; Vernasco *et al*. 2019; Ryder *et al*. 2020). The amount of time a male spends in the net has a subtle, but significant, negative effect on his circulating testosterone (Vernasco et al. 2019). Therefore, we used video monitoring to determine the duration of time that each bird was in the mist-net prior to blood sampling, so that it could be accounted for in further analyses (mean = 17.5 minutes, SD = 10.3 minutes, range = 1–72 minutes). Following removal of a male from the mist-net, a small blood sample (< 125uL) was taken from the brachial vein and stored on ice prior to being centrifuged at 10,000 rpm for 5 min, as described in previous studies (Ryder *et al*. 2011b; Vernasco *et al*. 2019; Ryder *et al*. 2020). Plasma volume was measured to the nearest 0.25 ul and stored in 0.75 ml of 100% ethanol (Goymann *et al*. 2007). Testosterone was double extracted from the plasma using dichloromethane. Following extraction, a direct radioimmunoassay was used to measure the total plasma androgen concentration (ng/ml) adjusted by the extraction efficiency and plasma volume of each sample (Eikenaar *et al*. 2011; Ryder *et al*. 2011b). Hormone assays were conducted annually, and the detection limits were 0.12, 0.08, and 0.09 ng/ml for 2015-16, 2016-17 and 2017-18, respectively; any sample that fell below the assay-specific limit of detection was assigned that limit as its testosterone concentration as a most conservative estimate. The extraction efficiency for all samples was between 62-73%, and the intra-assay coefficients of variation were 6.6%, 11.6%, and 9.2% for 2015-16, 2016-17 and 2017-18, respectively; the inter-assay coefficient of variation was 19.5%.

### Behavioral Recording

We used an automated data-logging system to monitor male-male interactions on the display territories of the leks (Ryder *et al*. 2012; Dakin & Ryder 2018; Ryder *et al*. 2020). The territories on these leks are specifically used for male-male coordinated social displays (as described in Schwartz & Snow 1978). At the beginning of each field season, male manakins were outfitted with coded nano-tags (NTQB-2, Lotek Wireless; 0.35 g). The tags transmitted a unique VHF signal ping once per 20 s for three months. In total, 296 tag deployments were performed on 180 individuals (mean 1.7 field seasons per male ± SD 0.7), 178 of whom also had hormone data (mean number of hormone samples per male = 3 ± SD 1.5). Approximately 10 days (± SD 7) after tagging and sampling was completed at a given lek, a proximity data-logger (SRX-DL800, Lotek Wireless) was deployed within each display territory at the lek to record tagged males within a detection radius of 30 m (a distance that corresponds to the typical diameter of a manakin display territory; Heindl 2002; Dakin & Ryder 2018).

Proximity recording sessions ran from 06:00 to 16:00 for ∼6 consecutive days (± SD 1 day) and were performed ∼3 times per field season at a given lek. Occasionally, the length of a recording session was extended due to extenuating circumstances such as inclement weather. The recording sessions were scheduled to be distributed evenly throughout each field season at each lek, to minimize any confounding seasonal effects. Each recording session represents an observation of the social network at a given lek. In total, we conducted 86 recording sessions (29,760 data-logger hours) representing repeated measures of the social activity at 11 leks during three field seasons (see Fig. S1 in the supplement for additional details on the sampling regime).

Prior to data-logger deployment, each territory was also observed on non-recording days to identify the territory-holder based on his color-bands, following previous studies (Ryder *et al*. 2008, 2009). These status assignments were subsequently verified in the proximity data.

### Data Processing and Statistical Analysis

All computational and statistical analyses were performed in R (R Core Team 2018). Network illustrations were made using the igraph package (Csardi *et al*. 2018).

### Social Interactions

Social networks were constructed by first defining interactions between two males that occurred on the display territories. To do this, a computational algorithm was used to identify joint detections, wherein two males were located at the same display territory within a pre-defined spatial and temporal threshold (Ryder *et al*. 2008, 2012; Dakin & Ryder 2018). For the temporal threshold, the two males had to occur < 45 s apart. This temporal threshold was chosen to allow for the fact that each tag pinged with a 20 second pulse rate, such that overlapping individuals could have up to a 40 s gap between their respective pings. For the spatial threshold, the two males had to have a difference in received signal strength values (Δ RSSI) < 10. This threshold corresponded to a typical distance between 0 to 5 m apart in a ground-truthing experiment (Dakin & Ryder 2018). Hence, according to our definition, a social interaction is initiated only after males come within this approximate spatial threshold. We chose this spatial threshold because it is close enough to permit visual and acoustic contact during typical social behaviors (such as those described in Schwartz & Snow 1978). After completing our study, we also performed a sensitivity analysis to verify that our main results were robust to alternative spatial threshold definitions (see supplement for details).

Any repeated co-occurrence of the two males within 5 minutes was considered to be part of the same social interaction, but after a gap of ≥ 5 minutes, it was considered to be a new interaction between those two males. The average duration of social interactions defined by this method was 5.1 min ± SD 13.3, 8.8 min ± SD 20.1, and 7.0 min ± SD 20.5 in the three respective field seasons, further indicating that these were sustained social interactions, rather than random encounters.

In total, we identified 36,885 social interactions over the three field seasons of study. These interactions were used to define a weighted social network for each lek recording session. The nodes in the network were the individual males, and the edges were weighted by the number of social interactions between each pair of males. An earlier validation study compared social interactions that were detected by the proximity system with those that were directly observed for 11 males (Ryder *et al*. 2012), and confirmed that all of the interactions detected by the proximity system were also directly observed.

### Null Model Validation of the Social Networks

Two broad classes of methods have been described for building social networks: (i) networks that are built based purely on the proximity of animals unrelated to their behavioral context (“gambit of the group”), and (ii) networks that are built based on specific behavioral criteria that are directly observed (Franks *et al*. 2010; Croft *et al*. 2011; Farine 2015). Although the interactions in our study were not directly observed, they were recorded at specialized display perches that have a known function in male-male social interactions (Schwartz & Snow 1978; Ryder *et al*. 2011a). Hence, they do not qualify as the gambit of the group. To further demonstrate this point, we performed a pre-network permutation of the raw data to determine whether the observed network edges occurred more often than expected by chance (Farine 2017). This analysis was highly conservative in that it preserved key features of the data including male visit rates to specific territories within each recording session and lek; it is described in detail in the supplement and Fig. S2. The results demonstrated that 95% of the observed network edges had a greater edge weight than expected under stringent permutation conditions. Moreover, these preferred edges had edge weights that were 10-fold to 50-fold greater than expected by chance under these stringent conditions (Fig. S2). This provides an additional validation to our approach, because it indicates that the observed network edges were nearly always preferred, even relative to other possible interactions within the same lek. These results are also consistent with a previous validation study indicating that these methods capture male-male coalition partnerships that are directly observed (Ryder *et al*. 2012).

Although pre-network data permutations are sometimes used to derive adjusted association indices, or to prune networks prior to further analysis, we did not take this approach for several reasons. First, the statistical rarity of a relationship in our system does not *a priori* define the importance of any one social interaction, especially given that all of the interactions occurred on display territories with specialized function. A single interaction between two rarely interacting males may have been highly consequential (e.g., if one of those males had a highly influential hormone-behavioral phenotype). Conversely, a single interaction between two frequently interacting males may have been relatively unimportant. It would be unwarranted to assume that partnership rarity indicates the importance of any single interaction in this context. Second, and perhaps more importantly, the goal of this study was a comparison across multiple networks. Some networks are genuinely less preferential, and more random, than others. If we modified the network edges based on permutation-based indices, it would disproportionately prune the truly random networks, introducing a source of bias that would be contrary to our main goal. Hence, all further analyses are based on networks where the edge weights are given by the observed number of interactions, as this method is most appropriate for our study system and aims. Below, we also describe a separate node-label permutation that provided an additional check on our final statistical analyses.

### Social Specialization

To quantify social specialization, we sought a metric that would capture the extent to which a network was partitioned into exclusive social relationships (as opposed to a network made up of non-specific or non-exclusive partnerships). To do this, we used a network metric of specialization that is commonly used in community ecology called H_2_’ (Blüthgen *et al*. 2006). An advantage of H_2_’ is that it is standardized against a theoretical maximum, based on the overall activity levels of different nodes and Shannon entropy (Blüthgen *et al*. 2006); this makes it possible to compare the extent of specialization across different bipartite networks in a standardized way. To apply this metric to our manakin data, we converted each lek’s social network into its bipartite adjacency matrix (Fig. 1A), with floaters along one axis, and territory-holders on the other, and then calculated social specialization as H_2_’ using the bipartite package (Dormann *et al*. 2019). Higher values of specialization indicate that the network is well-partitioned (i.e., made up of exclusive relationships), as illustrated in Fig. 1A. We chose to focus on floater-territorial specialization because these two social classes are well-defined and floater-territorial partnerships tend to be the most common in this species (Fig. S3; see also Ryder *et al*. 2011a).

Our measure social specialization at the network level, H_2_’, can also be related to the exclusivity of social partnerships at the individual level, as used in (Sih *et al*. 2009; Edenbrow *et al*. 2011; Dakin & Ryder 2018). All else being equal, a highly specialized network is expected to be made up of individuals who are relatively more exclusive and/or more important towards their partners, *sensu* (Sih *et al*. 2009; Dakin & Ryder 2018). Because H_2_’ has the properties described in the previous paragraph, it is more appropriate as the network-level metric of social specialization. Note that it is possible, and perhaps even common, for the edge weights in a highly specialized network to be relatively invariant if the network is not fully connected, as shown in the example manakin networks in Fig. 1A. Hence, specialization based on H_2_’ is not in principle related to the network-average coefficient of variation of each individual’s edge weights (Maldonado-Chaparro *et al*. 2018).

### Network Stability

We define network stability as the average persistence of network edges through time (Dakin & Ryder 2020). To quantify the stability of manakin networks, we compared each lek’s social network from a given recording session to its subsequent recording session within the same field season (Fig. 1B). Network stability was then calculated as the number of male-male partnerships (binary network edges) shared by both time points divided by the number of partnerships at either time point (Dakin & Ryder 2020). Higher values of stability indicate greater persistence of social relationships within the network, independent of any changes in the representation of particular males (nodes) (Poisot *et al*. 2012). To focus this measure on the persistence of strong relationships, we computed network stability of partnerships that occurred at least six times following (Dakin & Ryder 2020). The threshold of six was chosen because it corresponds to an average rate of one social interaction per day in our data. We conducted an additional sensitivity analysis to verify that alternative thresholds for the stability calculation (greater or less than six) did not change our main results (see supplement for details). Previous work using this metric of stability has shown that the wire-tailed manakin social networks are far more stable than expected by chance (Dakin & Ryder 2020).

### Behavioral Assortment

Assortment refers to the extent to which individuals associate with similar partners (Fig. 1C); it can be due to partner choice (homophily), shared environments, or the social transmission of behavior (Dakin & Ryder 2018). Assortment was quantified using Newman’s assortativity index, which is a correlation coefficient for the statistical association among linked nodes within a network. It ranges from –1 (a negative association), through 0 (no association), and up to +1 (a positive association). To quantify the assortment of social behaviors, we first determined the daily frequency of two behaviors for each male: his number of social interactions per day (strength), and his number of unique social partnerships per day (degree). Strength and degree are both repeatable measures of a male’s social behavior in our study population (Dakin & Ryder 2018). We used the average log-transformed values of each male’s strength and degree within the recording session, and then calculated the assortativity coefficient for the entire social network using the algorithm for weighted networks in the assortnet package (Farine 2016).

Because assortativity values for strength and degree were highly correlated (Pearson’s r = 0.78, p < 0.0001, n = 86 networks), we took the average of these two values as the measure of overall behavioral assortment within the social network. Note that we used log-transformed values of strength and degree because these two variables are strongly positively skewed (Dakin & Ryder 2018; Ryder *et al*. 2020), and assortativity is based on a Pearson’s correlation. Finally, we also computed the assortativity of the two discrete status classes (floater and territory-holder), to ensure that our analysis of behavioral assortment was not solely driven by status-assortment.

### Collective Testosterone

To understand how hormones might predict network properties, we derived a measure of collective testosterone of each social network. This was based on the hormonal trait that was the best predictor of social behavior in our previous study, referred to as “mean testosterone” (Ryder *et al*. 2020). A male’s mean testosterone is his average residual circulating testosterone. It is calculated using a linear regression of log-transformed testosterone to statistically account for the effects of field season, Julian date, time of day when captured, and duration of restraint, all of which may influence point estimates of a male’s baseline hormone level (Vernasco *et al*. 2019). Hence, mean testosterone represents a standardized measure of a male’s circulating testosterone, independent of his capture conditions (Fig. 2A). Next, to determine collective testosterone, we took the average mean testosterone for each social network, weighted by the interaction frequency (strength) of the males within the network (Fig. 2B). Collective testosterone is thus a group-level characteristic that is weighted towards the males that made the greatest contribution to group social structure. In other words, networks with low collective testosterone are made up of mostly low-testosterone individuals, whereas networks with high collective testosterone are made up of mostly high-testosterone individuals. We calculated collective testosterone separately for each status class, because the effects of hormones on social behavior are status-dependent (Eisenegger *et al*. 2011; Boksem *et al*. 2013; Ryder *et al*. 2020).

### Statistical Analysis

To evaluate the hypothesis that collective testosterone predicts network properties, we analyzed mixed-effects models of the social network properties in the lme4 package (Bates *et al*. 2018). The three response variables were social specialization, network stability, and behavioral assortment (Fig. 1). We used Akaike’s Information Criterion (AIC) to compare four candidate models for each response variable, as follows: (1) collective testosterone of territory-holders + collective testosterone of floaters; (2) collective testosterone of territory-holders; (3) collective testosterone of floaters, and (4) no testosterone predictors. All of the models included additional fixed effects to account for field season (a categorical variable with three levels), the average Julian date of the recording session, the average number of recorded hours per territory, and the size of the social network (number of individuals), as well as a random effect of lek to account for repeated measures.

Model comparison was performed on models fit with maximum likelihood, and the best-fit models were re-fit using restricted estimation of maximum likelihood (REML) to derive parameter estimates (Zuur *et al*. 2009). We used the lmerTest package to compute p-values for parameter estimates in the mixed-effects models based on Satterthwaite’s method (Kuznetsova *et al*. 2018). We also report Nakagawa and Schielzeth’s R^2^_LMM_ values as an estimate of effect size (Nakagawa & Schielzeth 2013). We verified that all models met the assumptions of linear regression analyses. First, we checked that the Pearson residuals met the assumption of normality. We visually inspected the partial residual plots for each fixed and random effect, to confirm that there were no outliers or departures from the assumption of homoscedasticity. To check for multicollinearity, we used the performance package (Lüdecke *et al*. 2019) to calculate variance inflation factors (VIFs), and we verified that all VIFs were between 1 – 1.8.

### Sample Sizes and Exclusions

In two field seasons (2016-17 and 2017-18), we performed an experiment as part of a separate study to test the influence of transiently-elevated testosterone on individuals (n = 5 individuals in 2016-17 and n = 4 in 2017-18; Ryder *et al*. 2020). The results of that experiment demonstrated that elevated testosterone caused a temporary decrease in the frequency and the number of social partnerships in the altered males (Ryder *et al*. 2020). It is important to note that this experiment was not designed to test emergent properties at the network level, because it was conducted on a limited scale with only one or two individuals temporarily altered within each lek. We therefore excluded the 6 post-manipulation networks from the main analysis in this study. We verified that when we included these manipulated leks, all of our main conclusions were unchanged.

After excluding the 6 post-manipulation observations, 80 of the original 86 recording sessions remained. Table 1 summarizes the sample sizes for the network-level analyses. In 10 cases, specialization could not be calculated because the bipartite network did not have sufficient data to determine H_2_’. Stability could not be calculated in 26 cases, when either the recording session occurred at the end of a field season, or when there were insufficient partnerships that met the criteria for the stability calculation.

### Node-label Permutation Analysis

To evaluate the possibility that our results could be influenced by other properties of the leks that were independent of hormonal traits, we also performed a statistical permutation of the post-network data (Farine 2017). The purpose of this analysis was to verify that the results were driven by (and sensitive to) the relative contribution and social position of different males. To do this, we performed a statistical permutation that randomized the node labels (male IDs) within each of the social networks, retaining network topology, and leaving each male’s testosterone traits unchanged. Hence, this analysis preserved which males were present in which recording session, but it randomized the relative position and contribution of each male. After generating 1,000 of these node-label permutation datasets, we recalculated specialization, stability, and assortment, and then refit the top models from our mixed-model analysis. We then compared the slope estimates from the observed data to those derived from 1,000 node-label permutations. As a one-sided p-value, we calculated the proportion of slope estimates from the node-label permutations that were more negative than the corresponding estimate from the observed data.

## Results

The lek social networks had 14 males on average, but there was considerable variation among leks ranging from 3 to 43 individuals (Fig. S1). Additional descriptive statistics are provided in Table 1. Although network size has a lognormal distribution (Fig. S1), all other emergent properties of the social structure were approximately normally distributed (Fig. S4).

We found that the collective testosterone of territorial males could predict all three emergent properties of the social networks (specialization, stability, and assortment). The leks with greater representation of high-testosterone territorial males were less specialized, less stable, and more negatively assorted (Fig. 3; all p ≤0.03 in mixed-effects models). The slope coefficients for these three relationships were also all greater than expected under a node-label permutation (inset panels, Fig. 3; all p < 0.02). Model selection results for the observed data indicated considerable uncertainty in the best-fit model for each of the three network properties analyzed (Table S1). However, the collective testosterone of territory-holders was a significant predictor in all of the best-supported models (Table S2). In contrast, the collective testosterone of floater males was not a significant predictor of network properties in any of the best-fit models (Table S2).

**Figure 3.**
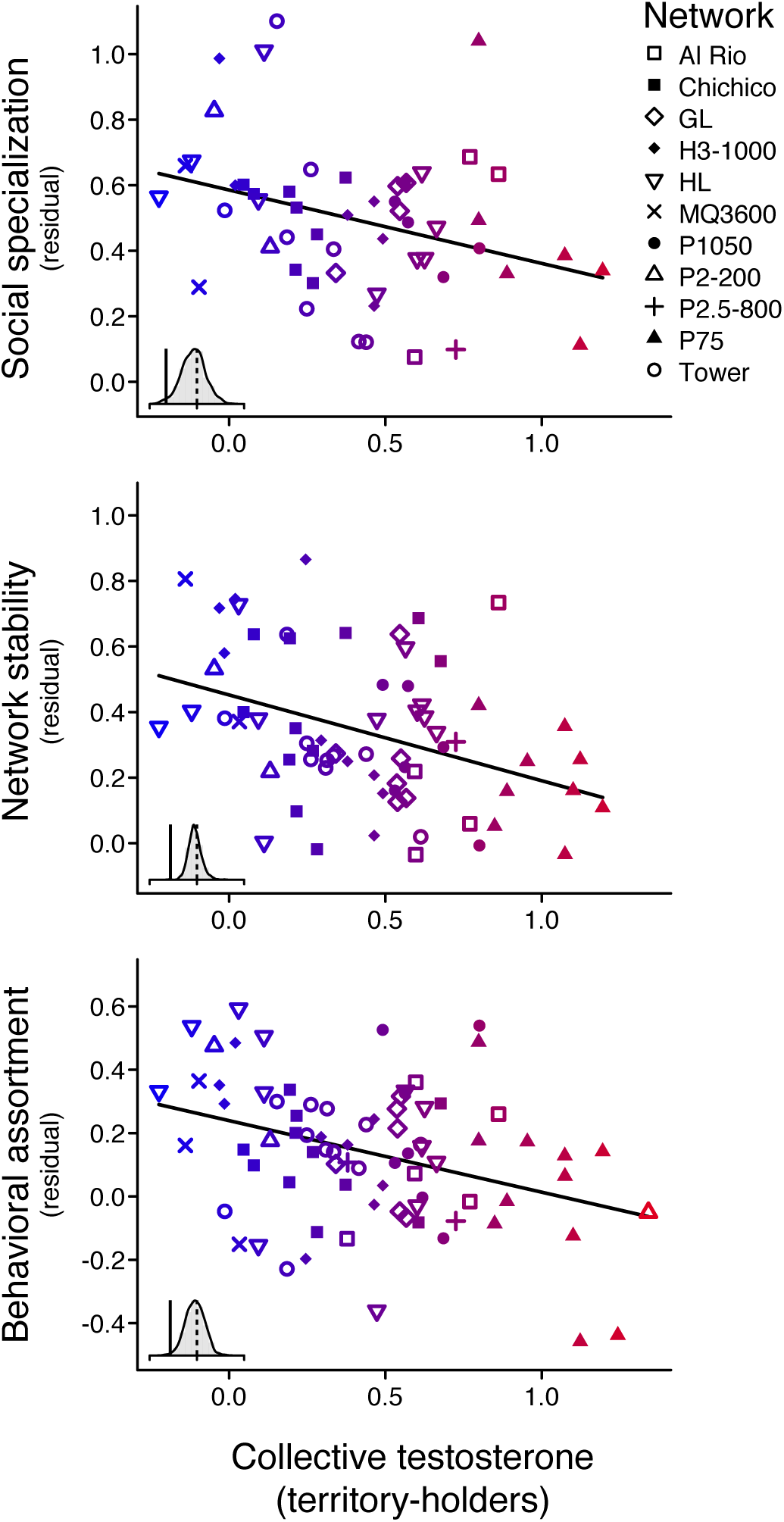
Collective testosterone predicts the structure of the social network. The networks with higher-testosterone dominant males were less specialized, less stable over time, and had more negative behavioral assortment. Each plot shows the partial residuals from a statistical analysis that also accounts for field season, the average Julian date of the recording session, the number of recording hours, network size, and lek identity. Because floater and territorial males can differ in behavior, the analysis of behavioral assortment also accounted for status assortment within each network. Different symbols are used to indicate repeated measures of 11 different leks, colored according to the collective testosterone scale in Fig. 2. Inset panels show the results of node-label permutation tests. In each case, the slope in the best-fit model (solid vertical line) is significantly more negative than expected based on the distribution of permuted estimates (grey distribution, dotted line is at 0). See Tables S1-S2 for additional data.

We did not detect any significant effects of Julian date on network properties within our study period, but we did observe some significant year-to-year differences (e.g., social specialization and behavioral assortment in Table S2). Assortment was the only property that was significantly related to network size and recording effort (Table S2). All else being equal, behavioral assortment was more positive in larger networks, and it was more negative in networks that had longer recording sessions.

R^2^_LMM(m)_ provides an estimate of the proportion of variance explained by the fixed effects in a model (Table S1). The R^2^_LMM(m)_ for assortment in the best-fit model was 0.52. This indicates that about 52% the variation in behavioral assortment could be explained by the combined associations with collective testosterone, network size, sampling effort, and annual variation (Table S2). The R^2^_LMM(m)_ for stability was 0.12, indicating that collective testosterone and the other predictors (network size, sampling effort, year) could explain about 12% the variation in that metric (Table S2). Finally, for specialization, the R^2^_LMM(m)_ indicated that the combined effects of testosterone and these other sources of variation together explained about 20% of the variation (Table S2).

## Discussion

The collective testosterone of the dominant, territory-holding males within a lek was associated with multiple emergent properties of the social network (Fig. 3). Variation in collective testosterone is a function of both the number of high-testosterone males and their frequency of social interactions. Our results indicate that the hormone-mediated behavior of these individuals may affect all three social network properties of specialization, stability and assortment. This indicates that the effects of testosterone on dominant males may mediate an extended phenotype with the power to shape social structure (Dawkins 1982). In contrast, the collective testosterone of floater males was not significantly associated with any of the emergent network properties (Table S2).

How can the relationship between hormone levels and network properties be explained in terms of individual mechanisms? Given that testosterone has antagonistic effects on the sociality of territorial males (Ryder *et al*. 2020; Vernasco *et al*. 2020), we hypothesize that the behavior of high-testosterone individuals can cause several features of the social organization to break down. Our previous results showed that high-testosterone dominant individuals have a reduced ability to attract and maintain social partners (Ryder *et al*. 2020). We propose that this weakening of social relationships may cause floater males to prospect elsewhere for new partners, negatively affecting both the stability and specialization of the lek network as a whole. Likewise, high-testosterone dominant males may inhibit the processes of social contagion, reciprocity, and/or behavioral matching that can cause positive behavioral assortment (Dakin & Ryder 2018). Recent studies have found that in other social animals, sparse and specialized social networks can be associated with fitness benefits (Stroeymeyt *et al*. 2018); hence, a breakdown to this organization may incur costs (Maldonado-Chaparro *et al*. 2018). Testing our proposed mechanism for the link between hormone-mediated behavior and network dynamics will require direct observation of the individual social behaviors that occur within dyads, and how these behaviors change through time. Because our current data cannot assess the fine-scale valence of social interactions, further studies are needed that combine direct observation with high-throughput data on social network dynamics.

We included several additional parameters in our analyses to account for social network size and sampling effort. Although it was not one of our main hypotheses, we noted that network size was positively associated with behavioral assortment (Table S2). In other words, males were more likely to associate with behaviorally similar partners in larger leks, whereas they were more likely to associate with dissimilar partners in smaller leks. Effects of group size on assortment have been noted in a few other studies, although the form of this relationship varies (Griffiths & Magurran 1997; Ilmarinen *et al*. 2017; McDonald *et al*. 2017). In manakins and other lekking systems, larger leks are known to have heightened display activity and higher female visitation rates (Lank & Smith 1992; Durães *et al*. 2009). This raises the possibility that social facilitation and heightened activity may be associated with increased homophily and/or behavioral matching (either through partner choice, contagion, or reciprocity). Another plausible explanation is that smaller social groups may constrain these behavioral processes, by making it more difficult to find or match a suitable partner. This hypothesis could be explored in experiments on captive systems and simulation models.

Behavioral assortment was also more positive in networks that were recorded for less time in our study (in other words, leks where males associated with behaviorally similar partners tended to be recorded for fewer days). Recording time was designed to be approximately even among leks (Fig. S1), with the most common reason for an extended recording duration being inclement weather that extended the recording session. Therefore, we speculate that inclement weather may have reduced the amount of behavioral matching (assortment) while also affecting recording time. Although we did not collect data on weather at each lek as this was not our main goal, the question of how inclement weather affects group-level social dynamics is an interesting one that merits further study.

Conducting experimental tests of causation at the level of whole social networks remains a major challenge in ecology (Pinter-Wollman *et al*. 2014; James *et al*. 2009). Although our node-label permutation analysis provides evidence that our results are not merely due to structural differences among networks, we cannot rule out the possibility that high-testosterone individuals chose to participate in certain networks due to other factors that may also influence the emergent properties of the network (e.g., environmental quality and/or female activity). It is important to note that in this system, as in many other wild animals, controlled experimental manipulations of the broader social network structure are not yet possible (Zyphur *et al*. 2009; Akinola *et al*. 2016). Nevertheless, our data here indicate that the increased prevalence of dominant, high-testosterone individuals can predict changes social dynamics and subsequent higher-order network structure. These findings establish that hormone-behavior relationships are not limited to one individual, but instead hormones have population-level consequences (McClintock 1981; Robinson 1992).

Increasing evidence demonstrates that the structure and stability of social networks is often associated with benefits–and costs–during foraging, breeding, and disease outbreaks (Silk *et al*. 2010; Maldonado-Chaparro *et al*. 2018; Riehl & Strong 2018; Stroeymeyt *et al*. 2018). Given the widespread influence of steroid hormones on social interactions across vertebrates (Adkins-Regan 2005), we expect that collective testosterone will be broadly associated with social network properties in other systems. The direction of these effects may depend on the context of the behavioral interactions that form the social network, and whether these interactions are primarily cooperative or competitive in nature. Taken together with previous studies, we propose that testosterone-mediated behavior can alter social network dynamics in ways that often impede the evolution of stable social relationships and cooperation.

## Acknowledgments

We thank Camilo Alfonso, Brian Evans, David and Consuelo Romo, Kelly Swing, Diego Mosquera, Gabriela Vinueza, and the Tiputini Biodiversity Station of the Universidad San Francisco de Quito. Funding was provided by the National Science Foundation (NSF) IOS 1353085 and the Smithsonian Migratory Bird Center.

## Author Contributions

RD, ITM, BMH, and TBR designed the study. ITM, BMH, BJV, and TBR collected the data. RD, ITM, and TBR analyzed the data. RD and TBR wrote the manuscript. All authors edited the manuscript.

## Data Availability

All data and R scripts necessary to reproduce this study are available for download at: https://figshare.com/s/13a311662fee686fa4f3

## Supporting Information

Supplemental text, Figures S1-S5, and Table S1-S3 in the attached PDF

