## Supplementary Information for "Testosterone-mediated behavior shapes the emergent properties of social networks"

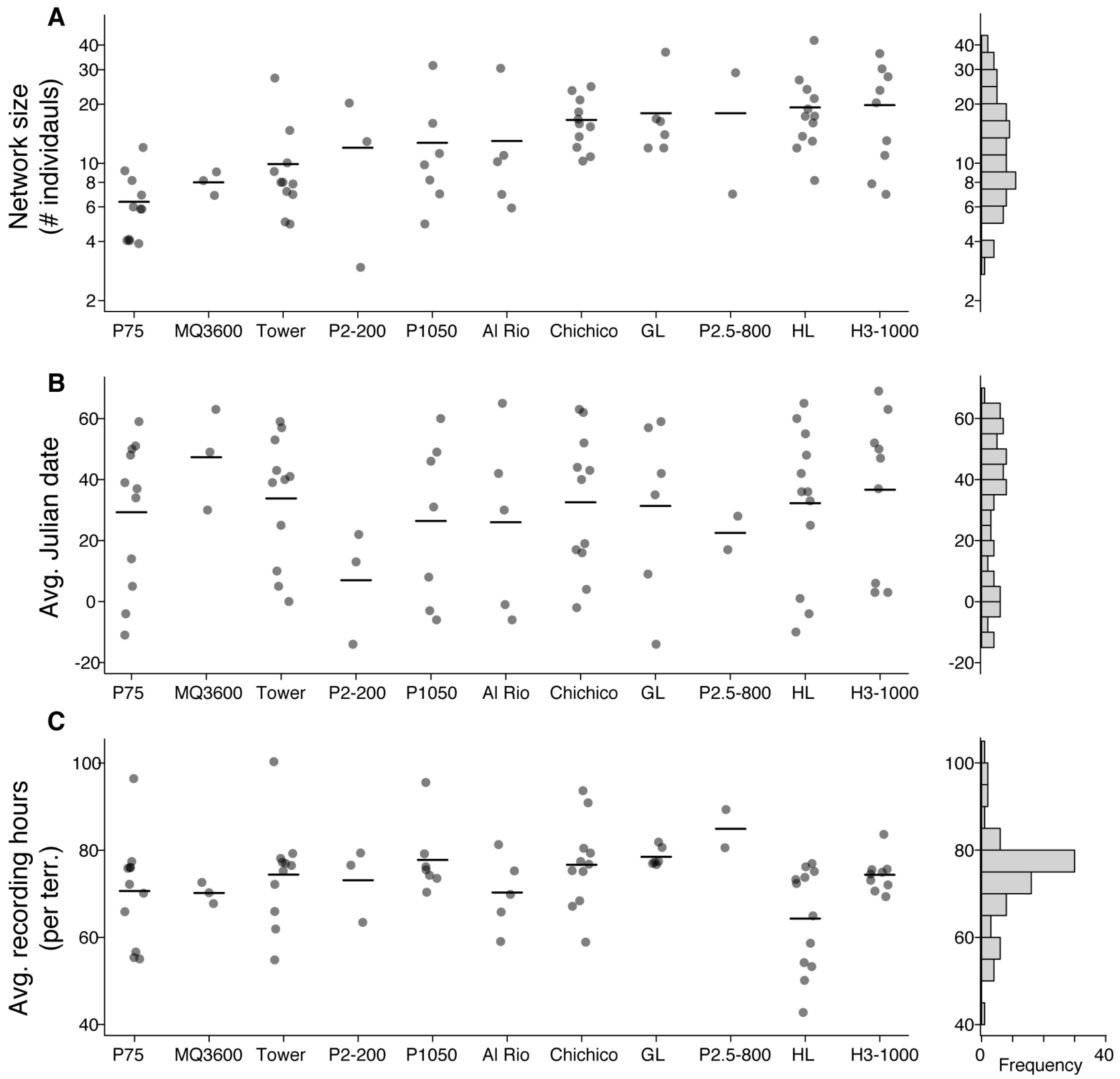


**Figure S1. Network size and sampling regimes.** Each of the 11 leks is plotted along the x-axis. Data are shown for (A) social network size, (B) the average Julian date, and (C) the average number of recording hours per territory, for each recording session. The horizontal lines display the means. Note that network size in (A) is plotted on a log scale, and the 11 leks are ordered by average size throughout the figure. Random jitter is also used throughout the figure to show datapoints that would otherwise overlap.

***Pre-network Permutation Analysis***

The goal of the pre-network permutation analysis was to validate that most of the network edges occurred more often than expected in a permutation of the raw data. To do this, we permuted the timing of each individual's raw visitation bouts to specific display territories within each recording session. The first step was to define and label the individual visit events, which we defined as a series of consecutive pings by the same male on a given territory, with no gaps > 90 s. The threshold of 90 s was chosen because it was twice the length of the 45 s threshold used to define social interactions. If a male had a 90 s gap with no pings at a given territory, the next ping he made on that same territory was considered to be a new visit event. The raw data contained over 1.04 million visits defined by these criteria.

After giving each visit its own unique ID label, we performed a permutation to shuffle the date and starting time of the first ping of each visit within each territory. The new date and time of the first ping were chosen randomly from the observed data for that specific recording session and territory. Any subsequent ping times within the visit were then recalculated, such that the duration and relative ping times of the original visit were preserved. This permutation was both complete (i.e., all visit events were permuted in each step), and it was also highly constrained to preserve key features of the data. Specifically, it preserved the number of times that each male visited each territory within each recording session, the variation among males in their overall activity rates, and the typical time of day of the visits within each territory and recording session.

After the permutation was performed, we re-ran the algorithm described in the main text to identify social interactions in the permuted data (i.e., any events wherein two males were at the same territory within 45 s and $\Delta$ RSSI < 10). This algorithm performs a chronological search and is computationally intensive, so we were limited to performing only 100 full iterations of this analysis. We therefore examined correlations among the first 100 iterated results to confirm that they were generally consistent. We focused on correlations in the edge weights for the 2,998 session-partnerships that also occurred in the real data, since these were most critical to further interpretations. The correlation in permuted edge weights for those 2,998 partnerships was strong and highly consistent (mean $r = 0.60$ [95% CI = 0.60 – 0.61] for 100 randomly-selected correlation pairs; each Pearson's correlation had $n = 2{,}998$). Moreover, the correlation between the average permuted edge weight, and any one iteration, was also remarkably strong (mean $r = 0.78$ [95% CI = 0.78–0.78] for 100 pairs; each Pearson's correlation $n = 2{,}998$). Given the high consistency in these initial checks, we infer that there would be little information gained by performing additional iterations of the pre-network data permutation.

The overall results of the pre-network data permutation analysis are shown in Fig. S2 below. Most notably, the number of social interactions that occurred in the permuted datasets was lower than that observed in the actual dataset by a factor of seven (mean = 5,154 per permutation dataset [95% CI = 5,138 – 5,170], as compared to 36,885 in the actual observed dataset). Furthermore, 95% of the observed edge weights were greater than expected relative to the permutation. These results highlight how remarkably specific the real interactions are, even compared to a highly conservative raw data permutation.

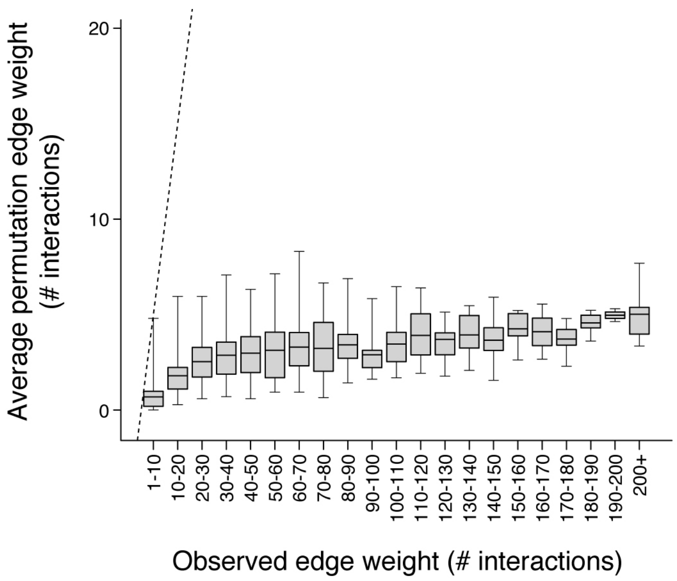


**Figure S2. Results of the pre-network permutation analysis.** Average permutation edge weight is plotted in relation to actual edge weight (n = 2,998 session-partnerships that actually occurred). Each boxplot shows the mean, IQR, and range for the permutation data. The dotted line denotes the region where y = x (i.e., where the permutation occurrence is equal to actual occurrence). Out of 2,998 partnerships that actually occurred, 2,840 (95%) occurred more often than expected in the pre-network permutation. This indicates that the observed partnerships were nearly always preferred.

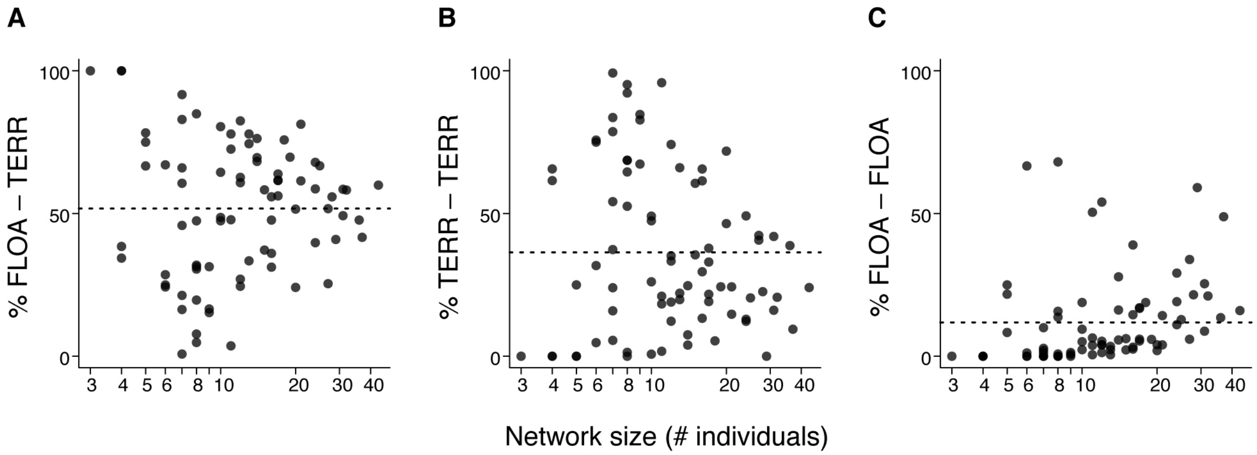


**Figure S3. The composition of social interactions in the manakin social networks.** (A) The most common type of social interaction was between a floater and a territorial male. These interactions made up 52% of observed social networks, on average, although there was considerable variation among networks as shown in (A). (B) The second most common type of interaction involved two territory-holding males. This type of interaction made up 36% of the observed social networks, on average. (C) Interactions between two floater males were the least common, making up only 12% of the social networks, on average. The horizontal dotted lines show the means. This figure is limited to unmanipulated leks from our main analysis (n = 80 recording sessions).

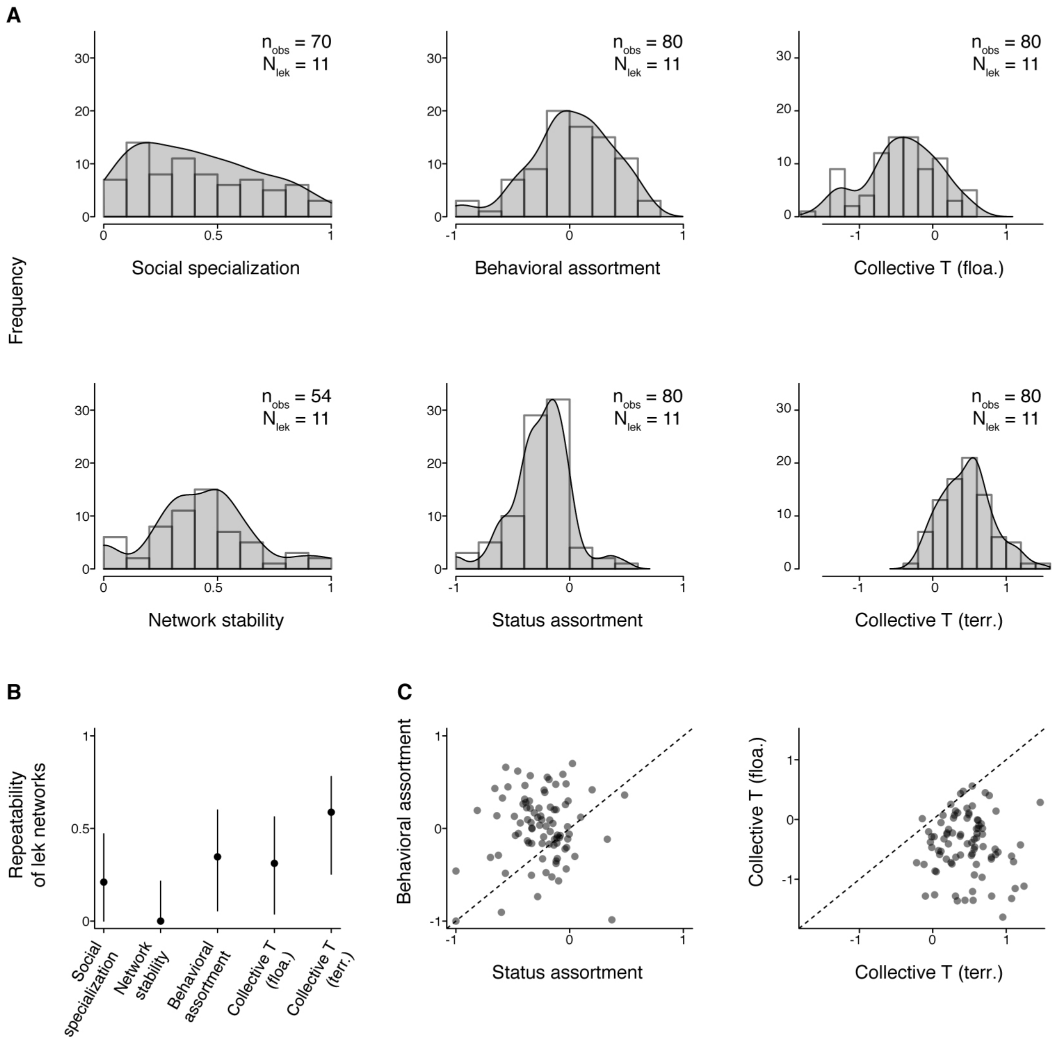


**Figure S4. Properties of the manakin social networks.** (A) Histograms and smoothed probability density curves are shown for six social network properties. The sample sizes are provided within each panel. (B) Repeatability of network properties, calculated as the proportion of total variance attributed to differences among the 11 leks. Error bars show 95% confidence intervals determined by parametric bootstrapping. (C) Scatterplots showing the association between similar metrics. The first scatterplot shows the association between behavioral assortment and status assortment of a network. The second scatterplot shows the association between the collective testosterone of floater males and territorial males. The dotted lines in (C) denote the region where y = x. Neither of the correlations in (C) is statistically significant (all $p > 0.10$).

**Table S1. Candidate models of specialization, stability, and assortment.** All models include a random effect of lek, as well as fixed effects to account for field season, Julian date within the field season, the number of individuals in the social network, and the average number of recording hours per territory. Models of behavioral assortment include an additional fixed effect for the assortment of the two status classes within the network. The evidence ratio provides the likelihood that the best-fit model (ranked first) is a better fit to the data as compared to the *i^th^* ranked model. The sample size is lower for specialization and stability because these two metrics could not be calculated for all networks (see main text for details). The marginal r-squared statistic, $R^2_{LMM(m)}$, provides an estimate of the proportion of total variance explained by the fixed effects in the best-fit model. The conditional r-squared statistic, $R^2_{LMM(c)}$, provides an estimate of the proportion of total variance explained by both the fixed effects and the random effect of lek.

| Model set | Candidate model, fixed effects | ΔAIC | Akaike weight | Evidence ratio (best-fit vs. model i) | $R^2_{LMM(m)}$ (best-fit model) | $R^2_{LMM(c)}$ (best-fit model) |
|---|---|---|---|---|---|---|
| Social | 2. Testosterone terr. | 0 | 0.60 | – | 0.20 | 0.33 |
| specialization | 1. Testosterone terr. + Testosterone floa. | 1.34 | 0.31 | 2.0 | | |
| n = 70 | 4. (No Testosterone) | 4.41 | 0.07 | 9.1 | | |
| | 3. Testosterone floa. | 6.23 | 0.03 | 22.2 | | |
| Network | 2. Testosterone terr. | 0 | 0.58 | – | 0.12 | 0.13 |
| stability | 1. Testosterone terr. + Testosterone floa. | 1.57 | 0.27 | 2.2 | | |
| n = 54 | 4. (No Testosterone) | 3.44 | 0.11 | 5.6 | | |
| | 3. Testosterone floa. | 5.15 | 0.05 | 13.0 | | |
| Behavioral | 1. Testosterone terr. + Testosterone floa. | 0 | 0.49 | – | 0.52 | 0.58 |
| assortment | 2. Testosterone terr. | 0.31 | 0.42 | 1.2 | | |
| n = 80 | 3. Testosterone floa. | 4.74 | 0.05 | 10.7 | | |
| | 4. (No Testosterone) | 4.84 | 0.04 | 11.2 | | |

**Table S2. Best-fit models of specialization, stability, and assortment.** Slope estimates are provided from fixed effects in the best-fit models from Table S1. All p-values are two-sided tests, based on Satterthwaite's approximation for mixed-effects models in the lmerTest package. For each response variable, we provide details of the two best-supported models, because the AIC values indicated similar support for top two models of each response variable. Notably, the conclusions regarding collective testosterone are identical when comparing the top two models for each response variable.

| Response variable | Fixed effects | Estimate (SE) | t-value | p-value |
|---|---|---|---|---|
| Social specialization n = 70 | Field season | | | |
| | **2016-17 vs. 2015-16** | **–0.16 (0.08)** | **–2.11** | **0.04** |
| | 2017-18 vs. 2015-16 | –0.08 (0.08) | –1.03 | 0.31 |
| | 2017-18 vs. 2016-17 | 0.08 (0.07) | 1.11 | 0.27 |
| | Julian date | 0.002 (0.001) | 1.64 | 0.11 |
| | Network size (log) | –0.008 (0.07) | –0.12 | 0.91 |
| | Recording hours | 0.001 (0.003) | 0.40 | 0.69 |
| | **Collective T (terr.)** | **–0.26 (0.11)** | **–2.49** | **0.02** |
| | Field season | | | |
| | **2016-17 vs. 2015-16** | **–0.17 (0.08)** | **–2.18** | **0.03** |
| | 2017-18 vs. 2015-16 | –0.06 (0.08) | –0.69 | 0.49 |
| | 2017-18 vs. 2016-17 | 0.11 (0.08) | 1.33 | 0.19 |
| | Julian date | 0.002 (0.001) | 1.66 | 0.10 |
| | Network size (log) | –0.007 (0.07) | –0.10 | 0.92 |
| | Recording hours | 0.001 (0.003) | 0.38 | 0.71 |
| | Collective T (floa.) | 0.05 (0.07) | 0.69 | 0.50 |
| | **Collective T (terr.)** | **–0.26 (0.10)** | **–2.54** | **0.02** |
| Network stability n = 54 | Field season | | | |
| | 2016-17 vs. 2015-16 | 0.09 (0.08) | 1.03 | 0.32 |
| | 2017-18 vs. 2015-16 | –0.006 (0.08) | –0.07 | 0.94 |
| | 2017-18 vs. 2016-17 | –0.11 (0.08) | –1.33 | 0.19 |
| | Julian date | –0.001 (0.002) | –0.84 | 0.41 |
| | Network size (log) | 0.002 (0.07) | 0.02 | 0.98 |
| | Recording hours | –0.004 (0.004) | –0.99 | 0.33 |
| | **Collective T (terr.)** | **–0.22 (0.10)** | **–2.23** | **0.03** |
| | Field season | | | |
| | 2016-17 vs. 2015-16 | 0.09 (0.09) | 1.08 | 0.29 |
| | 2017-18 vs. 2015-16 | 0.02 (0.09) | 0.20 | 0.84 |
| | 2017-18 vs. 2016-17 | –0.07 (0.08) | –0.86 | 0.40 |
| | Julian date | –0.002 (0.002) | –0.93 | 0.36 |
| | Network size (log) | –0.01 (0.07) | –0.17 | 0.86 |
| | Recording hours | –0.003 (0.004) | –0.91 | 0.37 |
| | Collective T (floa.) | 0.05 (0.08) | 0.61 | 0.55 |
| | **Collective T (terr.)** | **–0.23 (0.10)** | **–2.24** | **0.03** |
| Behavioral assortment n = 80 | Field season | | | |
| | 2016-17 vs. 2015-16 | –0.003 (0.07) | –0.05 | 0.96 |
| | **2017-18 vs. 2015-16** | **0.16 (0.07)** | **2.11** | **0.04** |
| | **2017-18 vs. 2016-17** | **0.16 (0.07)** | **2.21** | **0.03** |
| | Julian date | 0.002 (0.001) | 1.70 | 0.09 |
| | **Network size (log)** | **0.36 (0.06)** | **6.46** | **< 0.0001** |
| | **Recording hours** | **–0.007 (0.003)** | **–2.67** | **0.009** |

| | | | |
|---|---|---|---|
| Status assortment | 0.08 (0.11) | 0.73 | 0.47 |
| Collective T (floa.) | 0.08 (0.06) | 1.38 | 0.17 |
| **Collective T (terr.)** | **–0.22 (0.09)** | **–2.57** | **0.009** |
| Field season | | | |
| 2016-17 vs. 2015-16 | –0.001 (0.07) | 0.008 | 0.99 |
| 2017-18 vs. 2015-16 | 0.12 (0.07) | 1.68 | 0.10 |
| 2017-18 vs. 2016-17 | 0.12 (0.07) | 1.77 | 0.08 |
| Julian date | 0.002 (0.001) | 1.76 | 0.08 |
| **Network size (log)** | **0.36 (0.06)** | **6.35** | **< 0.0001** |
| **Recording hours** | **–0.007 (0.003)** | **–2.58** | **0.01** |
| Status assortment | 0.07 (0.12) | 0.64 | 0.53 |
| **Collective T (terr.)** | **–0.22 (0.09)** | **–2.50** | **0.02** |

***Sensitivity Analysis***

We used two sensitivity analyses to check the robustness of our main results to parameter values used for processing the social network data.

The first sensitivity analysis focused on the spatial threshold used to define social interactions from the proximity data streams. Based on ground-truthing experiments, our main analysis used a threshold of < Δ10 RSSI units as indicative of a socially interacting dyad (Δ RSSI is a measure of the differential signal strength of two tags pinging the receiver at a given time). The threshold of Δ10 for social interactions was chosen based on a ground-truthing experiment showing that tags that were in the same location (i.e., less than 1 m apart), and tags that were within 5 m of each other, typically had signal differentials of less than Δ10 RSSI units. To determine whether alternative thresholds would affect our main conclusions about the relationship between testosterone and social structure, we re-processed all of the raw data using a series of alternative threshold values for this criterion: < Δ6, < Δ7, < Δ8, < Δ9, < Δ11, and < Δ12 RSSI units, respectively (smaller Δ values = closer tags, and a more stringent spatial definition). In each case, we redefined the social networks from the proximity data, recalculated all of the properties of the social networks including collective testosterone, and then repeated the statistical analyses of how social network properties relate to collective testosterone as described in the main text.

Our second sensitivity analysis focused on the metric of social network stability. This network property was calculated from networks that had been filtered to remove the weakest edges. The purpose of the filtering step was to ensure that stability would be based on strong partnerships, and not easily biased by rarely interacting dyads. In our original analysis, we used a filter of < 6 interactions per recording session, because this threshold corresponded to a rate of one social interaction per day in our dataset. In other words, network stability was based on strong dyads defined as occurring at least once per day, on average. To determine whether alternative filter thresholds would affect our conclusions, we re-calculated network stability using threshold values of < 3, < 4, < 5, < 7, < 8, and < 9, respectively, and then repeated the statistical analysis of network stability in relation to collective testosterone as described in the main text.

The results of all sensitivity analyses are summarized in Table S3 and Figure S5 below. Although these alternative parameters affected the absolute values of metrics (e.g., the number of social interactions in Table S3), they did not alter any of our main conclusions regarding the negative relationships between collective testosterone and emergent properties of the social networks. Overall, these results demonstrate that the relative values of social network properties, and the relationships between them, are robust to a wide range of parameter values during the initial data processing.

**Table S3. Sensitivity analyses of the model selection results.** We checked whether alternative data processing parameters would affect the results of the main statistical analysis in Tables S1-S2. ΔRSSI is the spatial criterion used to define social interactions from the raw data streams. The stability filter is the edge weight criterion used to define strong vs. rarely interacting dyads. In all cases, the best-fit model from the statistical analysis in Tables S1-S2 was the same as that found using the original data-processing parameters.

| Parameter | Value | Total # of social interactions | Best-fit model same as original analysis? | | |
|---|---|---|---|---|---|
| | | | Specialization | Stability | Assortment |
| **Δ RSSI** | < 10 (original) | 36,885 | – | – | – |
| *(smaller values* | < 6 | 30,689 | yes | yes | yes |
| *are more* | < 7 | 32,515 | yes | yes | yes |
| *stringent/* | < 8 | 34,100 | yes | yes | yes |
| *exclude more* | < 9 | 35,591 | yes | yes | yes |
| *cases)* | < 11 | 38,058 | yes | yes | yes |
| | < 12 | 39,133 | yes | yes | yes |
| **Stability filter** | < 6 (original) | – | – | – | – |
| *(larger values are* | < 3 | – | – | yes | – |
| *more stringent/* | < 4 | – | – | yes | – |
| *exclude more* | < 5 | – | – | yes | – |
| *edges)* | < 7 | – | – | yes | – |
| | < 8 | – | – | yes | – |
| | < 9 | – | – | yes | – |

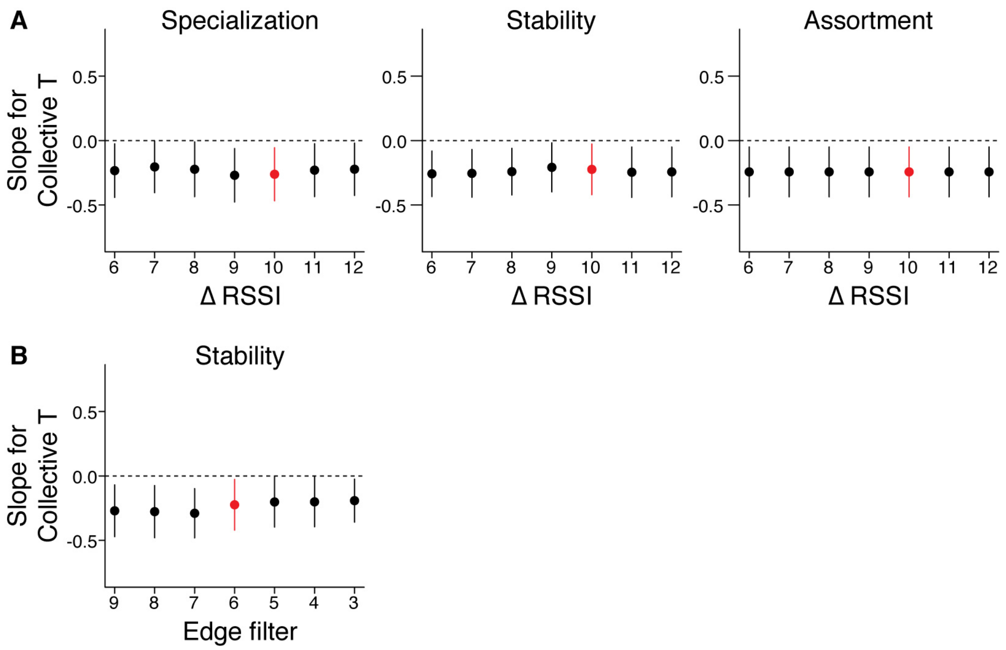


**Figure S5. Sensitivity analysis of slope estimates from the best-fit models.** Results are shown for alternative values of (A) Δ RSSI, the spatial criterion used to define social interactions, and (B) the edge weight criterion used to calculate stability. Each datapoint shows the slope coefficient from the best-fit model ± its 95% CI. The response variable is named above each plot. The original results from Table S2 are highlighted in red.
